# Spt6 is required for the fidelity of promoter selection

**DOI:** 10.1101/347575

**Authors:** Stephen M. Doris, James Chuang, Olga Viktorovskaya, Magdalena Murawska, Dan Spatt, L. Stirling Churchman, Fred Winston

## Abstract

Spt6 is a conserved factor that controls transcription and chromatin structure across the genome. Although Spt6 is viewed as an elongation factor, *spt6* mutations in *Saccharomyces cerevisiae* allow elevated levels of transcripts from within coding regions, suggesting that Spt6 also controls initiation. To address the requirements for Spt6 in transcription and chromatin structure, we have combined four genome-wide approaches. Our results demonstrate that Spt6 represses transcription initiation at thousands of intragenic promoters. We characterize these intragenic promoters, and find sequence features conserved with genic promoters. Finally, we show that Spt6 also regulates transcription initiation at most genic promoters and propose a model of initiation-site competition to account for this. Together, our results demonstrate that Spt6 controls the fidelity of transcription initiation throughout the genome and reveal the magnitude of the potential for expressing alternative genetic information via intragenic promoters.

## INTRODUCTION

While we once believed that transcription occurs primarily across coding regions, we now know that the transcriptional landscape is extraordinarily complicated, with transcription throughout the genome generating multiple classes of transcripts (Jensen et al., 2013; Pelechano, 2017). Regulation of these transcripts is exerted at several levels, including transcription initiation, elongation, termination, and RNA stability. The pervasive nature of transcription suggests that promoters are not only restricted to the 5’ ends of coding regions, but are widespread across the genome. How the cell defines and regulates initiation sites is therefore fundamental to gene expression.

Past genetic studies in yeast produced the unexpected finding that the specificity of transcription initiation may be controlled in part by transcription elongation factors, including histone chaperones and histone modification enzymes (Cheung et al., 2008; Hennig and Fischer, 2013; Kaplan et al., 2003). Therefore, transcription and co-transcriptional processes influence the permitted sites of transcription initiation within the genome. One factor that plays a critical role is Spt6, a conserved protein that directly interacts with RNA polymerase II (RNAPII) (Close et al., 2011; Diebold et al., 2010b; Liu et al., 2011; Sdano et al., 2017; Sun et al., 2010; Yoh et al., 2007), histones (Bortvin and Winston, 1996; McCullough et al., 2015), and the essential factor Spn1/Iws1 (Diebold et al., 2010a; Li et al., 2018; McDonald et al., 2010). Spt6 is believed to function primarily as an elongation factor based on its association with elongating RNAPII (Andrulis et al., 2000; Ivanovska et al., 2011; Kaplan et al., 2000; Mayer et al., 2010) and its ability to enhance elongation both in vitro (Endoh et al., 2004) and in vivo (Ardehali et al., 2009), although it has also been shown to regulate initiation in some cases (Adkins and Tyler, 2006; Ivanovska et al., 2011). During transcription, Spt6 regulates chromatin structure (Bortvin and Winston, 1996; DeGennaro et al., 2013; Ivanovska et al., 2011; Jeronimo et al., 2015; Kaplan et al., 2003; Perales et al., 2013; van Bakel et al., 2013) as well as histone modifications, including H3K36 methylation (Carrozza et al., 2005; Chu et al., 2006; Yoh et al., 2008; Youdell et al., 2008) and in some organisms, H3K4 and H3K27 methylation (Begum et al., 2012; Chen et al., 2012; DeGennaro et al., 2013; Wang et al., 2017; Wang et al., 2013). Substantial evidence suggests that a primary function of Spt6 is as a histone chaperone, required to reassemble nucleosomes in the wake of transcription (see (Duina, 2011) for a review).

Studies in the yeasts *S. cerevisiae* and *S. pombe* have shown that Spt6 controls transcription genome-wide (Cheung et al., 2008; DeGennaro et al., 2013; Kaplan et al., 2003; Pathak et al., 2018; Uwimana et al., 2017; van Bakel et al., 2013). In *spt6* mutants, the pattern of transcription dramatically changes, including altered sense transcription and increased levels of antisense transcription. Most notably, in *spt6* mutants there is extensive upregulation of cryptic or intragenic transcripts that appear to initiate from within protein-coding sequences (Cheung et al., 2008; DeGennaro et al., 2013; Kaplan et al., 2003; Uwimana et al., 2017). While many factors have been shown to control intragenic transcription, Spt6 is among the most broadly required for its regulation (Cheung et al., 2008).

In this work, we address longstanding issues regarding intragenic transcription and its regulation by Spt6 in *Saccharomyces cerevisiae*. Previous methods used to assay transcripts in *S. cerevisiae spt6* mutants (Northerns (Kaplan et al., 2003), tiled microarrays (Cheung et al., 2008), and RNA-seq (Uwimana et al., 2017)) could not distinguish whether intragenic transcripts were the result of new initiation or the result of RNA processing or decay. Microarrays and RNA-seq were also unable to detect intragenic transcripts from highly transcribed genes (Cheung et al., 2008; Lickwar et al., 2009). By comprehensively characterizing transcription initiation in wild-type and *spt6* strains with methods that directly assay initiation, we demonstrate that intragenic transcripts result from new initiation, and that Spt6 normally represses initiation from thousands of intragenic promoters. Furthermore, we characterize the chromatin structure and sequence features of intragenic promoters, and show that they share some sequence characteristics with canonical promoters at the 5’ ends of genes (hereafter referred to as genic promoters). Finally, we demonstrate that, contrary to previous beliefs, Spt6 widely controls transcription initiation from genic promoters and suggest that this is due to a competition between genic and intragenic promoters. Thus, Spt6 controls the fidelity of transcription initiation across the genome.

## RESULTS

### Spt6 regulates transcription initiation from intragenic promoters

To overcome the limitations of previous methods used to study transcription in *S. cerevisiae spt6* mutants, we adapted a transcription start site sequencing (TSS-seq) method (Arribere and Gilbert, 2013; Malabat et al., 2015) to identify the position of the RNA 5′-cap at single nucleotide resolution in both wild-type *S. cerevisiae* and in an *spt6* mutant. In the wild-type strain, TSS-seq was highly specific for reads mapping to annotated start sites, reproducibly identifying 4936 previously annotated TSSs (Malabat et al., 2015) (Figure 1A, Figure S1A, B). As TSS-seq measures the level of the 5’-ends of capped transcripts, we found a strong positive correlation between RNA levels measured by TSS-seq and RNA-seq for wild-type yeast (Uwimana et al., 2017) (Figure S1C). Thus, TSS-seq determines the positions of TSSs at high resolution and quantitatively measures the levels of capped RNAs.

**Figure 1.**
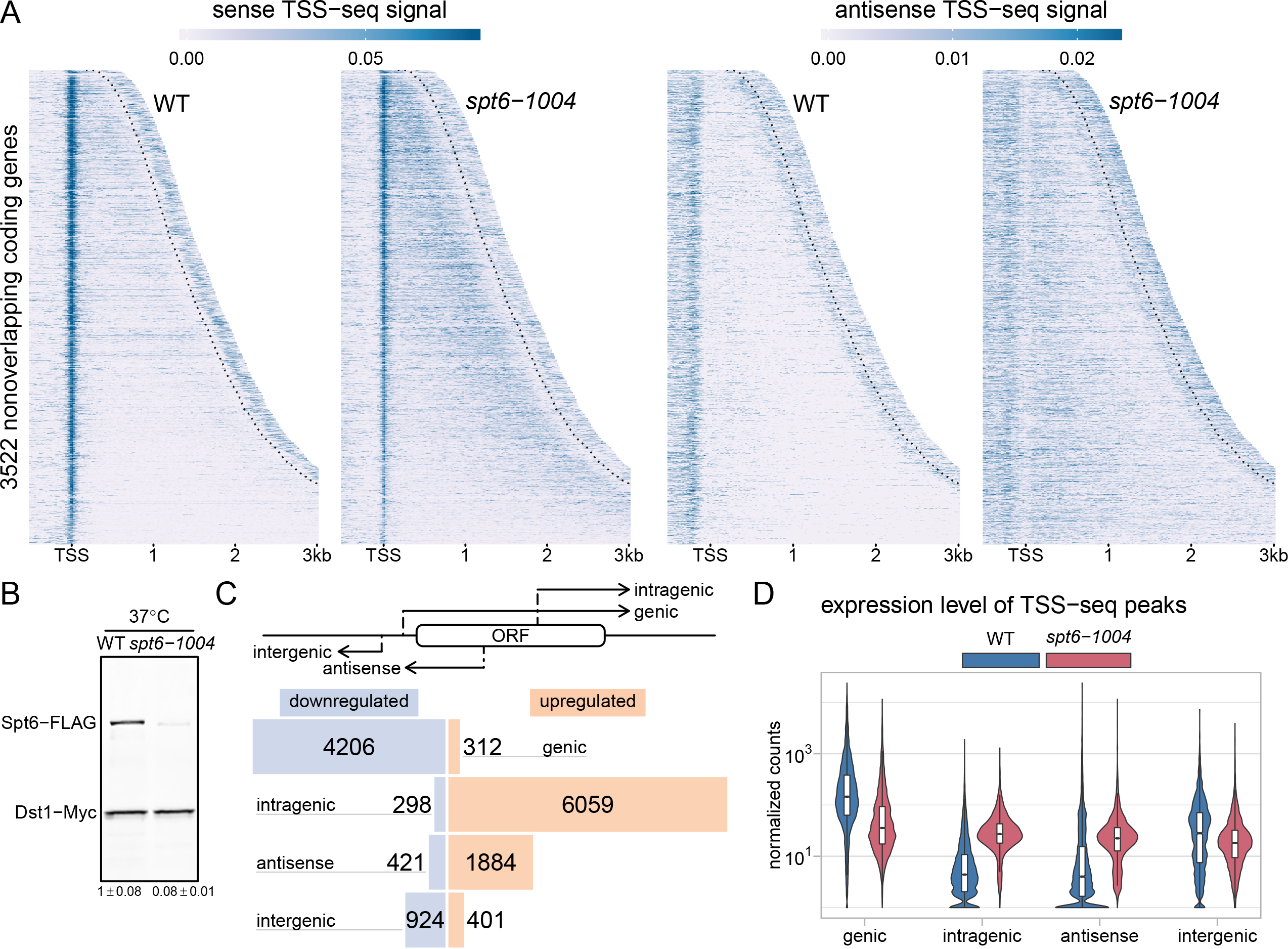
Spt6 is globally required for normal transcription initiation. **(A)** Heatmaps of sense and antisense TSS-seq signal in wild-type and *spt6-1004* cells, over 3522 non-overlapping genes aligned by wild-type genic TSS and sorted by length. Data are shown for each gene up to 300 nucleotides 3′ of the cleavage and polyadenylation site (CPS; indicated by the dotted line). Values are the mean of spike-in normalized coverage in non-overlapping 20 nucleotide bins, averaged over two replicates. Values above the 95th percentile are set to the 95th percentile for visualization. **(B)** Western blot showing the level of Spt6 protein in wild-type and *spt6-1004* strains after an 80-minute shift to 37°C. Protein levels were quantified using anti-FLAG antibody to detect Spt6 and anti-Myc to detect Dst1 from a spike-in strain (see Methods). The numbers below the blot show the average and standard deviation for three Westerns. **(C)** The diagram at the top illustrates the transcripts generated by the different classes of TSSs. The bar plot below shows the number of TSS-seq peaks differentially expressed in *spt6-1004* versus wild-type, classified by genomic region. Blue bars indicate downregulated peaks and orange bars indicate upregulated peaks. Differential expression was determined using DESeq2 (see Methods). **(D)** Violin plots showing the expression level distributions for different genomic classes of TSS-seq peaks in wild-type and *spt6-1004* strains. Values are the mean of counts from two replicates, normalized using an *S. pombe* spike-in (see Methods).

TSS-seq analysis of the *spt6-1004* mutant gave dramatically different results compared to wild type (Figure 1A). In our experiments, the *spt6-1004* mutation caused depletion of Spt6 to approximately 8% of wild-type levels after an 80-minute shift to the non-permissive temperature of 37°C (Figure 1B), although the cells were still viable (Kaplan et al., 2003). Under these conditions, we identified over 8,000 TSSs as significantly upregulated at least 1.5-fold in *spt6-1004* compared to the wild-type strain (Figure 1C). Approximately 6,000 of these TSSs are intragenic TSSs on the sense strand of a gene, although we also detect upregulated TSSs within annotated promoter regions, antisense intragenic (hereafter referred to as antisense), and in intergenic regions (Figure 1C). Our results show that intragenic TSSs are considerably more common than previously known and occur in approximately 60% of *S. cerevisiae* genes (Figure S1D). We note that sense strand intragenic TSSs tend to occur towards the 3’ ends of transcription units, while antisense TSSs tend to occur towards the 5’ ends (Figure 1A, S1E). We compared the set of genes we found that contain upregulated sense intragenic TSSs to the genes found by two previous genome-wide studies that identified sense intragenic transcripts in *spt6-1004* by microarrays (Cheung et al., 2008) and RNA-seq (Uwimana et al., 2017). We found considerable overlap between all three studies, though the use of TSS-seq allowed us to identify about 1,700 additional genes with at least one intragenic TSS (Figure S1F). Finally, we examined the expression levels of the different classes of transcripts as measured by TSS-seq and found that in the *spt6-1004* mutant, expression levels for all classes became more similar to one another (Figure 1E). Notably, our results revealed that the transcript levels are reduced from a majority of the genic TSSs, a result that we analyze in more detail in a later section. Taken together, our TSS-seq results demonstrate that the upregulation of thousands of capped and polyadenylated transcripts which occurs in an *spt6-1004* mutant is due to new transcription initiation, primarily within coding regions, and that this event is more widespread than previously known.

### Spt6 Controls the Localization of TFIIB

Given the dramatic changes in transcription initiation in an *spt6-1004* mutant, we wanted to assay transcription initiation using an independent approach, as well as to determine if intragenic promoters contain an RNAPII pre-initiation complex (PIC). Therefore, we measured genomic binding of TFIIB, an essential member of the RNAPII PIC, in wild-type and *spt6-1004* strains. To do this, we used ChIP-nexus (He et al., 2015), a modification of ChIP-exo (Rhee and Pugh, 2012), which measures the occupancy of a chromatin-bound protein at high resolution by exonuclease digesting the DNA up to the point of crosslinking and sequencing the position of the digested ends. We found that TFIIB binding patterns as measured by ChIP-nexus are reproducible (Figure S2A) and consistent with previous TFIIB ChIP-exo results at both the genome-wide scale and at TATA boxes (Figures S2B, S2C).

In the wild-type strain, the TFIIB ChIP-nexus signal was primarily localized upstream of previously annotated TSSs, as expected. Using the ChIP-seq peak-calling tool MACS2 (Zhang et al., 2008b), a TFIIB peak was found overlapping the window extending 200 base pairs upstream of 89% (4297/4917) of wild-type genic TSS-seq peaks. In contrast, in the *spt6-1004* mutant, the pattern of TFIIB binding across the genome was vastly altered, with TFIIB infiltrating coding regions in concordance with our TSS-seq results (Figure 2A, 2B). To test whether the increase in TFIIB binding over gene bodies might be caused by an increased level of TFIIB in the *spt6-1004* mutant, we measured TFIIB protein levels and found that they were actually reduced to approximately 70% of wild-type levels (Figure S2B). From these results we conclude that in the *spt6-1004* mutant, a more limited pool of TFIIB protein is much more widely associated across the genome than in wild type.

**Figure 2.**
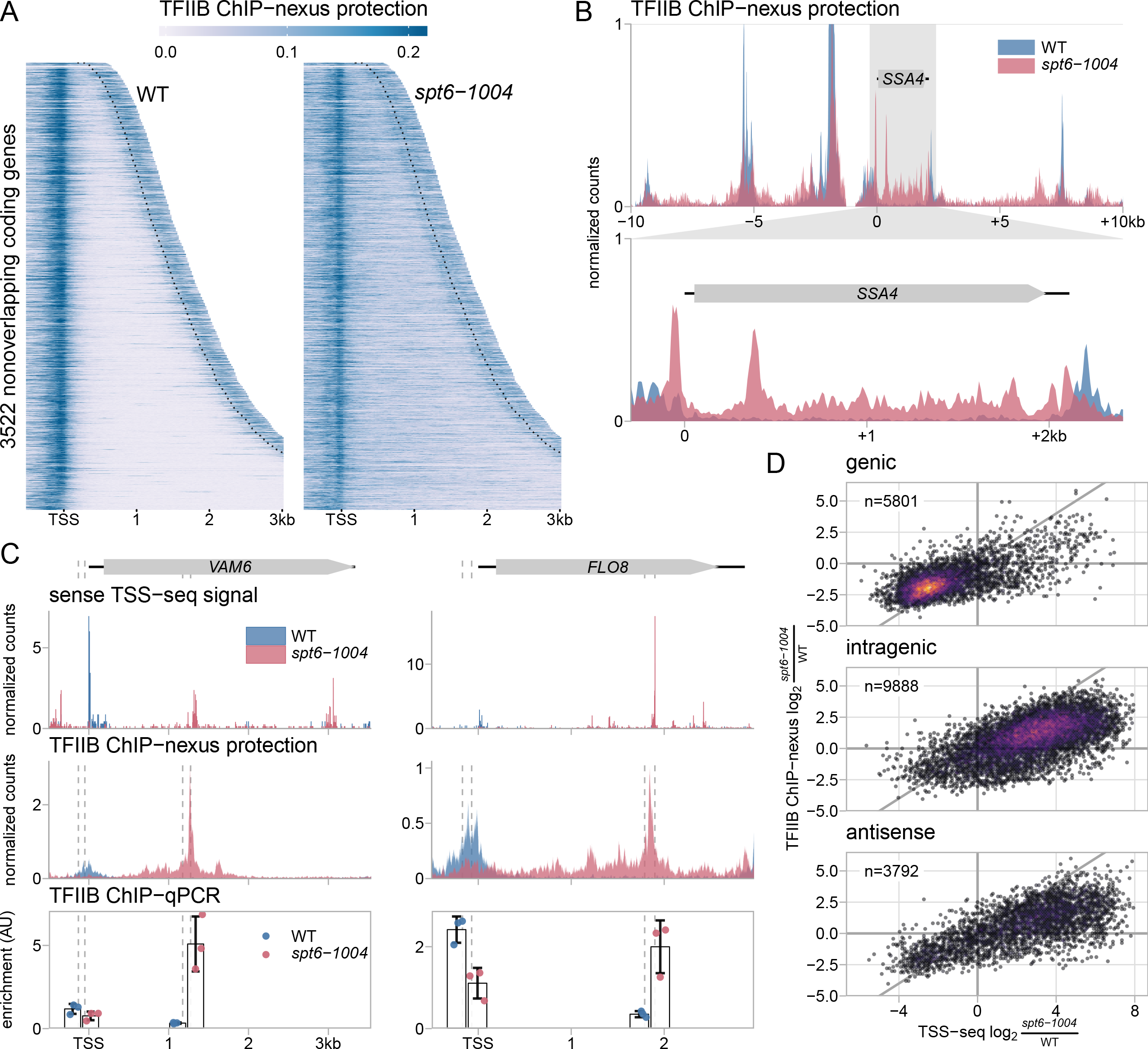
**Spt6 is required for** genome-wide localization of TFIIB. **(A)** Heatmaps of TFIIB binding as measured by ChIP-nexus in wild-type and *spt6-1004* strains, over the same regions shown in Figure 1A. The values are the mean of library-size normalized coverage in 20 basepair windows, averaged over two replicates. The position of the CPS is shown by the dotted lines. Values above the 90th percentile are set to the 90th percentile for visualization. **(B)** The upper panel shows TFIIB binding in wild-type and *spt6-1004* strains over 20 kb of chromosome II flanking the *SSA4* gene, as measured by TFIIB ChIP-nexus. The lower panel shows an expanded view of TFIIB binding over the *SSA4* gene. **(C)** TSS-seq, TFIIB ChIP-nexus, and TFIIB ChIP-qPCR measurements at the genic and intragenic promoters of the *VAM6* and *FLO8* genes in wild-type and *spt6-1004* strains. TSS-seq values are derived as described in Figure 1. ChIP-nexus values are as described above. ChIP-qPCR is normalized to amplification of a region of the *S. pombe pma1*^+^ gene used as a spike-in control. Vertical dashed lines represent the coordinates of qPCR amplicon boundaries. **(D)** Scatterplots of the fold-change in *spt6-1004* over wild-type strains, comparing TSS-seq and TFIIB ChIP-nexus. Each dot represents a TSS-seq peak paired with the window extending 200 nucleotides upstream of the TSS-seq peak summit for quantification of TFIIB ChIP-nexus signal. Fold-changes are regularized fold-change estimates from DESeq2, where size factors were determined from the *S. pombe* spike-in (TSS-seq) or from the *S. cerevisiae* counts (ChIP-nexus).

The altered binding pattern of TFIIB in *spt6-1004* (Figures 2A, 2B) made defining sites of intragenic initiation by TFIIB peak calling difficult. With the same parameters used to call peaks in the wild-type strain, MACS2 identified TFIIB peaks in *spt6-1004* upstream of 85% (4050/4763) of genic TSSs, but only identified TFIIB peaks upstream of 37.0% (2240/6059) of *spt6-1004* upregulated intragenic TSS-seq peaks. Two examples of these intragenic TFIIB peaks were verified by ChIP-qPCR of TFIIB (Figure 2C). Given the spreading-like nature of TFIIB association in many places in the *spt6-1004* mutant, it seemed plausible that there was an increased level of TFIIB upstream of the intragenic TSSs that are upregulated in *spt6-1004*, but that the nature of the TFIIB binding prevented a peak from being called. Therefore, we dispensed with TFIIB peak-calling and simply quantified the change in TFIIB signal in *spt6-1004* compared to wild type over the window extending 200 base pairs upstream of TSS-seq peaks. From this analysis, we found that the results from both assays were in agreement: 90.3% of genic promoters change in the same direction by both assays while approximately 81% of sense and antisense intragenic promoters change in the same direction (Figure 2D). We note that despite the challenge in calling intragenic TFIIB peaks, we did identify around 1500 intragenic TFIIB peaks that did not have a TSS-seq peak within 200 base pairs in either direction, which may represent intragenic initiation events not captured by TSS-seq, either due to non-productive initiation or transcript instability. Overall, the TFIIB ChIP-nexus results support our TSS-seq results and show that Spt6 controls TFIIB localization across the genome.

### Spt6 Controls Nascent Transcription on Both the Sense and Antisense Strands

As TSS-seq and ChIP-nexus measure steady-state levels of transcripts and PICs, respectively, we also performed native elongating transcript sequencing (NET-seq) (Churchman and Weissman, 2011) to measure the level and location of actively elongating RNAPII in wild-type and *spt6-1004* strains. Although NET-seq was unable to comprehensively provide information about intragenic transcription, it was able to provide other meaningful new information regarding the requirement for Spt6 in transcription. In wild-type cells, our NET-seq results were similar to those previously reported (Churchman and Weissman, 2011), with a high level of RNAPII density over approximately the first 750 bp of the sense strand of transcription units and a lower level further downstream. In contrast, in the *spt6-1004* mutant, we observed reduced levels of RNAPII over the 5’ region with a relative increase downstream (Figure 3A; S3A, S3B). The reduction in RNAPII density over the 5’ region provides independent evidence that genic transcription initiation is generally decreased in *spt6-1004*. The apparent increase in elongating RNAPII density over the 3’ regions of genes in *spt6-1004* is likely caused in part by a combination of intragenic initiation and a slower rate of elongation (Ardehali et al., 2009; Endoh et al., 2004).

**Figure 3.**
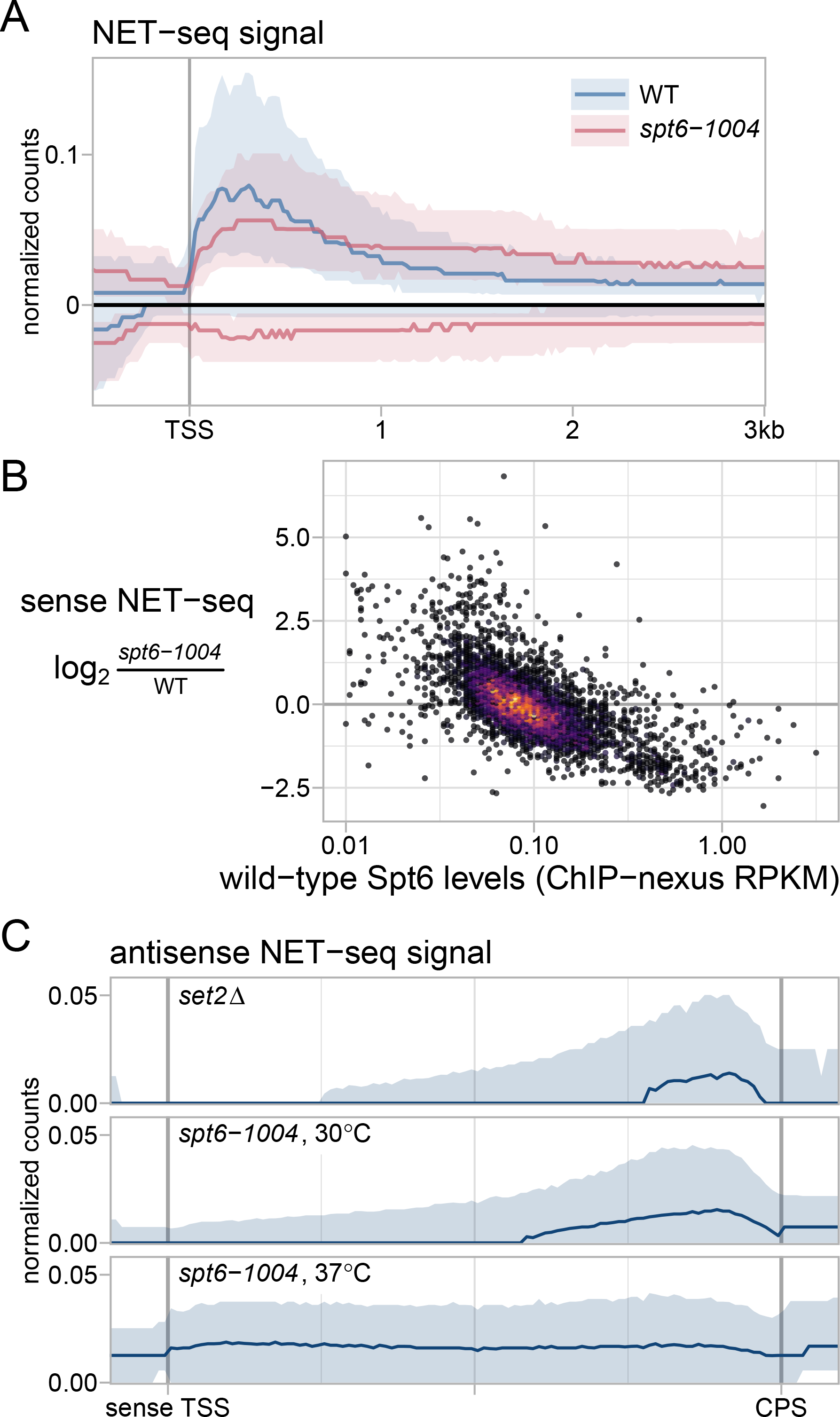
Spt6 is required for normal levels and distribution of elongating RNA polymerase II. **(A)** A metagene plot of the average sense and antisense NET-seq signals in wild-type and *spt6-1004* strains after a shift to 37°C, over 3522 nonoverlapping genes. Sense and antisense signals are depicted above and below the x-axis, respectively. The solid line and shadings represent the median and inter-quartile range, which are shown in order to give an idea of how the signal varies among the thousands of genes with diverse characteristics being represented in the plot. The values are the mean of library-size normalized coverage in nonoverlapping 20 nucleotide bins, averaged over two replicates. **(B)** A scatterplot of NET-seq fold-change in the *spt6-1004* mutant versus Spt6 occupancy in the wild-type strain as measured by Spt6 ChIP-nexus. Each dot represents a transcription unit for which sense NET-seq and Spt6 ChIP-nexus signals are summed over the entire length of the transcription unit. NET-seq fold-changes are regularized fold-change estimates from DESeq2. **(C)** Average antisense NET-seq signal in the *spt6-1004* strain at permissive (30°C) and nonpermissive (37°C) temperatures, compared to a *set2Δ* strain. The values are as in Figure 3A, and the solid line and shadings represent the median and inter-quartile range over 3522 nonoverlapping genes.

NET-seq also allowed us to test whether the level of Spt6 recruited to a gene corresponds to the degree of the requirement for Spt6 in active transcription. To do this, we performed ChIP-nexus of Spt6 in wild-type cells and compared that to the change in NET-seq signal in the *spt6-1004* mutant. From this analysis, we discovered a correlation between these two measurements: the genes with the greatest level of Spt6 in wild-type were those whose active sense-strand transcription was decreased the most in the *spt6-1004* mutant (Figure 3B). As there is a very strong correlation between the chromatin association of Spt6 and Rpb1 (Figure S3C; (DeGennaro et al., 2013; Ivanovska et al., 2011; Mayer et al., 2010; Perales et al., 2013)), this shows that the most highly transcribed genes are those that are most strongly dependent upon Spt6, in agreement with a recently published study (Pathak et al., 2018). These results independently support our TSS-seq and TFIIB ChIP-nexus results which suggested that transcription initiation from genic promoters is decreased in an *spt6-1004* mutant (Figures 1D, 2D), and further suggest that the degree of decrease correlates to the level of active transcription.

Our NET-seq results also revealed new information regarding Spt6 and antisense transcription. First, while our TSS-seq results suggested that most new antisense initiation in the *spt6-1004* mutant occurs towards the 5’ end of transcription units (Figure 1A), our NET-seq results showed antisense transcription to be elevated uniformly over the length of transcription units (Figure 3A, S3B). This difference likely results from high levels of antisense initiation from the 3’ UTR regions of genes (can be seen to right of the CPS line in Figure 1A; (Murray et al., 2012)). Second, as previous studies have demonstrated that *spt6-1004* mutants are defective for Set2-dependent H3K36 methylation (Carrozza et al., 2005; Chu et al., 2006; Youdell et al., 2008), and that *set2Δ* mutants also have elevated antisense transcription (Kim et al., 2016; Li et al., 2007; McDaniel et al., 2017; Venkatesh et al., 2016), we compared our NET-seq results for *spt6-1004* to previous NET-seq results for *set2Δ* (Churchman and Weissman, 2011). We included analysis of an *spt6-1004* mutant grown at 30°C, when Spt6 protein is still present, in addition to the same strain shifted to 37°C, when Spt6 protein is depleted. There is no detectable H3K36 methylation in the *spt6-1004* mutant at either temperature (data not shown). Our results (Figure 3C) show that *spt6-1004* grown at 30°C has a similar effect as *set2Δ* with respect to antisense transcription. However, after a shift to 37°C, the *spt6-1004* mutant has a greater derepression of antisense transcription than seen in *set2Δ.* These results suggest that the antisense effect in *spt6-1004* at 30°C is primarily due to loss of H3K36 methylation, while the effects seen after a shift to 37°C are additional *spt6-1004* specific effects, possibly due to changes in chromatin structure.

### Spt6 is Required for Normal Nucleosome Occupancy and Positioning

Several studies have shown that Spt6 is required for normal chromatin structure in *S. cerevisiae* (Bortvin and Winston, 1996; Ivanovska et al., 2011; Jeronimo et al., 2015; Kaplan et al., 2003; Perales et al., 2013; van Bakel et al., 2013). However, in order to correlate our TSS-seq results with high-resolution and quantitative analysis of chromatin structure, we performed MNase-seq to re-examine the requirement for Spt6 in maintaining chromatin structure. Our MNase-seq results from wild-type cells showed the expected signature over coding regions, including nucleosome-depleted regions 5’ of genes and a regularly phased pattern of nucleosomes over gene bodies (Figure 4A, S4A). In contrast, the pattern of nucleosome signal is drastically altered in the *spt6-1004* mutant, as previously observed (DeGennaro et al., 2013; van Bakel et al., 2013).

**Figure 4.**
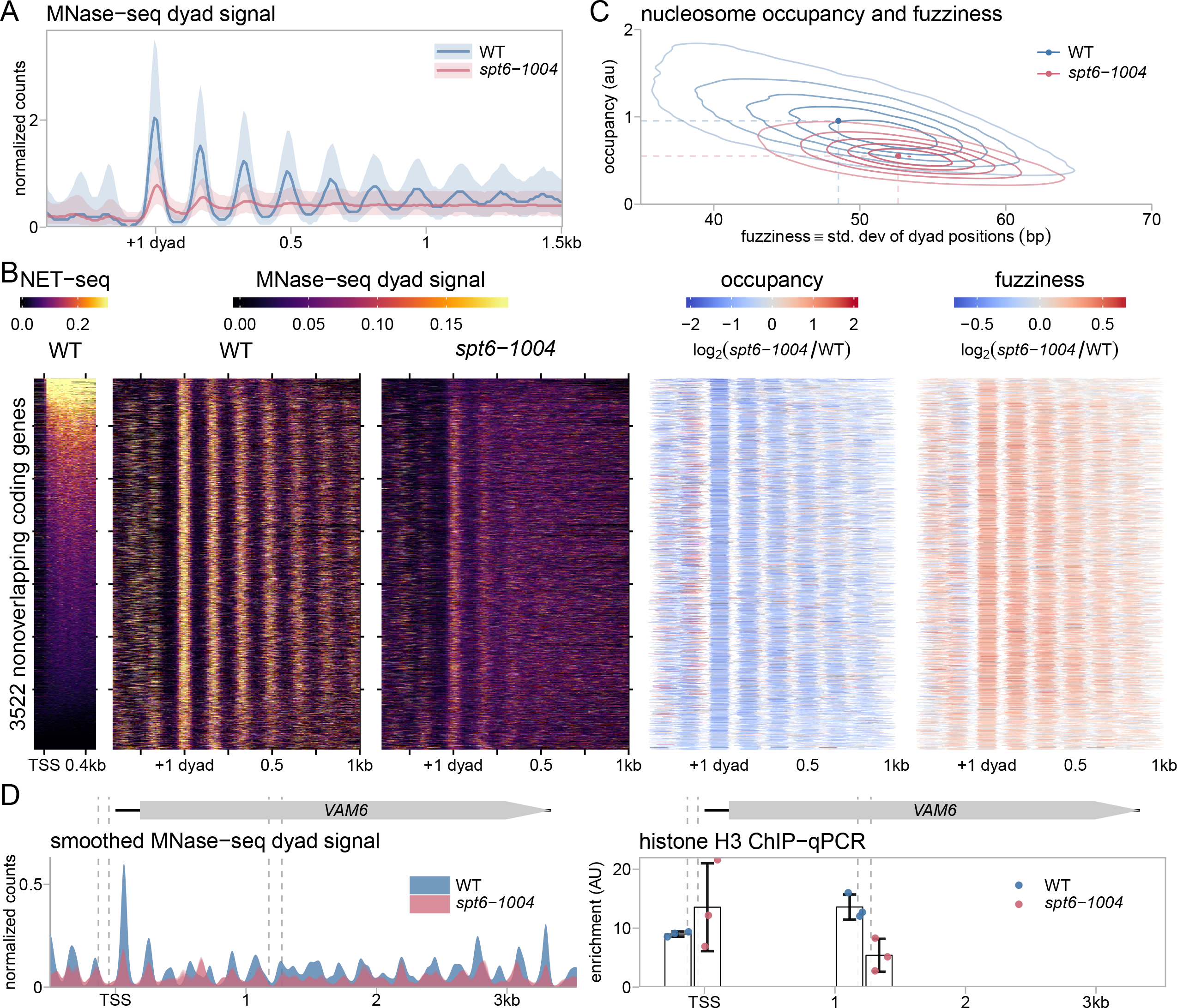
Genome-wide defects in chromatin structure in an *spt6-1004* mutant. **(A)** Average MNase-seq dyad signal in wild-type and *spt6-1004* strains, over 3522 nonoverlapping genes. The values are the mean of spike-in normalized coverage in nonoverlapping 20 nucleotide bins, averaged over two replicates *(spt6-1004)* or one experiment (wild-type). The solid line and shadings represent the median and inter-quartile range. **(B)** The leftmost panel shows the NET-seq signal in a window extending 500 nucleotides downstream of the TSS, sorted from top to bottom by the level of the signal. The second and third panels show heatmaps of the spike-in normalized MNase-seq dyad signal from wild type and *spt6-1004* strains over 3522 nonoverlapping coding genes aligned by wild-type +1 nucleosome dyad and sorted by total sense NET-seq signal. The last two panels show the spike-in normalized changes in nucleosome occupancy and fuzziness. The increased occupancy indicated just upstream of the +1 dyad is likely caused by nucleosomes occupying NDRs in the *spt6-1004* mutant. **(C)** A contour plot showing the global distribution of nucleosome occupancy and fuzziness in wild-type and *spt6-1004* strains. *(D)* MNase-seq and histone H3 ChIP-qPCR measurements of nucleosome signal at the *VAM6* gene in wild-type and *spt6-1004* strains. MNase-seq coverage is spike-in normalized dyad signal, smoothed using a Gaussian kernel with 20 bp standard deviation, and averaged by taking the mean of two replicates *(spt6-1004)* or one experiment (wild-type). Histone H3 ChIP-qPCR enrichment is normalized to amplification at the *S. pombe pma1*^+^ gene as a spike-in control. Vertical dashed lines represent the coordinates of the qPCR amplicon boundaries.

Differences in nucleosome signal are caused by different types of features, including occupancy and fuzziness (Chen et al., 2013). To determine the contribution of these to the altered nucleosome signal observed in *spt6-1004*, we quantified our MNase-seq data using DANPOS2 (Chen et al., 2013). In wild type, the population of nucleosomes varied greatly in occupancy and fuzziness, with more highly occupied nucleosomes tending to be less fuzzy (that is, more well positioned) (Figure 4B, 4C). In contrast, the distribution of nucleosomes in *spt6-1004* was more homogeneous, with a global decrease in occupancy and increase in fuzziness. To verity the decreased level of nucleosome occupancy, we performed histone H3 ChIP at three genes and found a lower level in the *spt6-1004* mutant compared to wild type, in agreement with previous results (Perales et al., 2013) (Figure 4D, S4C). This reduction may be caused, at least in part, by reduced expression of histone genes in *spt6* mutants (our TSS-seq data; (Compagnone-Post and Osley, 1996)). In summary, Spt6 plays a major role in determining nucleosome occupancy and positioning.

Previous work showed that genes with high levels of transcription show a relative decrease in nucleosome occupancy compared to genes with low levels of transcription (Shivaswamy et al., 2008). This trend is reflected in our wild-type MNase-seq data (Figure 4B). Furthermore, our previous work, based on the analysis of a much smaller number of genes, suggested that highly transcribed genes were most prone to nucleosome loss in an *spt6-1004* mutant (Ivanovska et al., 2011). However, from our new MNase-seq results, the severity of the changes in nucleosome signal in *spt6-1004* with respect to occupancy and fuzziness do not seem to depend on the transcription level (Figure 4B). We note that the weak nucleosome patterning observed in *spt6-1004* at highly transcribed genes compared to moderately transcribed genes is expected given that nucleosomes are already more disordered at highly transcribed gene in wild type (Figure 4B, S4B). These results, then, suggest that Spt6 controls chromatin structure genome-wide in a way that is independent of the level of transcription.

### Intragenic Promoters Have Some Sequence Characteristics of Canonical Promoters

Our TSS-seq analysis identified over 6,000 sense-strand intragenic TSSs that are derepressed in an *spt6-1004* mutant. To address how these promoters compare to canonical promoters at the 5’ ends of genes, we examined their chromatin structure and DNA sequence. Using the wild-type and *spt6-1004* MNase-seq data flanking the intragenic TSSs, we found that intragenic TSSs separated into two clusters that differed primarily by the phasing of the nucleosome array relative to the intragenic TSS (Figure 5A; see Methods). In wild-type chromatin, the intragenic TSSs from both clusters tended to occur at the border between regions of nucleosome enrichment and depletion, (Figure 5A), although nucleosome occupancy around these TSSs is modest compared to the occupancy adjacent to canonical promoters. This is likely due to the preference of sense-strand intragenic TSSs to occur towards the 3’ ends of transcription units. As expected, the average nucleosome signal around both clusters of intragenic TSSs is decreased in the *spt6-1004* mutant, consistent with derepression of transcription. In spite of the differences between the chromatin structure of the two clusters in wild-type strains, their expression levels in an *spt6-1004* mutant fall into a similar range (Figure 5B).

**Figure 5.**
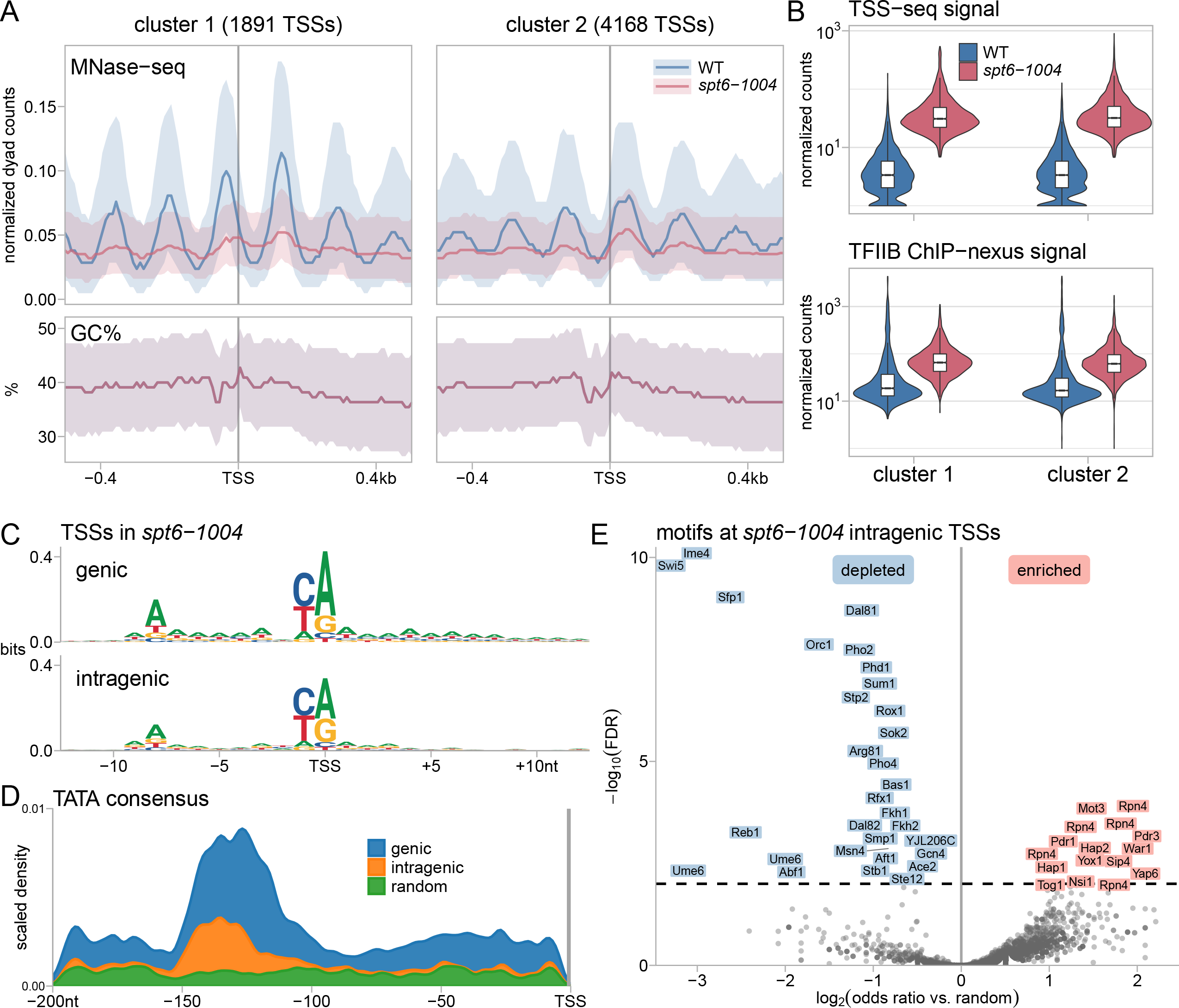
Chromatin structure and sequence features of intragenic promoters. **(A)** The average MNase-seq dyad signal and GC percentage for two clusters of intragenic TSSs that are upregulated in an *spt6-1004* mutant. The clusters were determined from the MNase-seq signal flanking the TSS (see Methods). **(B)** Violin plots showing the distributions of TSS-seq signal for the two clusters of intragenic TSSs that are upregulated in an *spt6-1004* mutant, and the distributions of their TFIIB ChIP-nexus signal in the window extending 200 nucleotides upstream of the TSS-seq peak. Counts are size factor normalized using the *S. pombe* spike-in (TSS-seq) or *S. cerevisiae* counts (TFIIB ChIP-nexus). **(C)** Sequence logos of the information content of TSS-seq reads overlapping genic and intragenic peaks in *spt6-1004* cells. **(D)** Scaled density of the TATA box upstream of TSSs. For each category, a Gaussian kernel density estimate of the positions of exact matches to the motif TATAWAWR is multiplied by the total number of TATA occurrences in the category and divided by the total number of regions in the category. **(E)** Volcano plot of motif enrichment and depletion upstream of intragenic TSSs upregulated in *spt6-1004*. Odds ratios and false discovery rate are determined by Fisher’s exact test, comparing to random locations in the genome. Some factors are shown more than one time because they had multiple motifs in the different databases that were searched.

Given that intragenic TSSs occur at specific sites, it seemed plausible that the alterations in chromatin structure might be a necessary factor for a potential intragenic promoter, but that chromatin structure alone was insufficient. Therefore, we also looked at the DNA sequence around the intragenic TSSs for particular sequences. First, as AT-rich sequences are unfavorable for nucleosomes and are often found in promoters (Iyer and Struhl, 1995; Kaplan et al., 2009; Tillo and Hughes, 2009; Zhang et al., 2009), we examined the GC content of the DNA sequence flanking intragenic TSSs and found a modest decrease in GC content just upstream of the TSSs in both clusters (Figure 5A, second row of panels). Second, we aligned the intragenic TSSs and discovered the same consensus initiation sequence, (A(A_rich_)_5_NPyA(A/T)NN(A_rich_)_6_), that was previously found for genic *S. cerevisiae* promoters (Malabat et al., 2015; Zhang and Dietrich, 2005) (Figure 5C). Third, we searched for TATA elements with perfect matches to the consensus sequence TATAWAWR (Basehoar et al., 2004). We found this consensus sequence at 10.7% of the regions upstream of *spt6-1004* sense-strand intragenic TSSs, compared to 23.7% for all genic TSSs and 8.8% over random sites in the genome. The TATA elements found upstream of intragenic TSSs tended to be located in the region 100 to 150 base pairs upstream of the TSS, the same region where TATA boxes upstream of genic TSSs are found (Figure 5D). When we limited our search to the top 1000 most significantly upregulated intragenic TSSs (out of 6059), the percentage of regions containing a TATA element increased to 15.4%. In summary, intragenic promoters are enriched for classes of sequence elements found at many genic promoters.

Finally, we quantified the enrichment or depletion of sequence-specific transcription factor binding site motifs upstream of intragenic TSSs and found many members of both classes (Figure 5E). The most enriched motifs are for transcription factors that are activated by cellular stresses (for example, Rpn4, Pdr1/3, and Mot3). This supports a previous observation that some intragenic promoters can be induced by stress (Cheung et al., 2008; McKnight et al., 2014; Tamarkin-Ben-Harush et al., 2017) (see Discussion). We also observed a significant depletion for multiple motifs, including those for Abf1 and Reb1, two factors that function in the establishment and/or maintenance of NDRs at many genic promoters (Badis et al., 2008; Kaplan et al., 2009; Lee et al., 2007; Tsankov et al., 2010; Yarragudi et al., 2007). The depletion for these motifs highlights the lack of a typical NDR for intragenic promoters.

### A General Requirement for Spt6 in Genic Promoter Function

Our TSS-seq data revealed the unexpected finding that Spt6 is required for normal expression levels from most genic promoters. Out of 5,274 genes, 3,857 (73.1%) were downregulated in the *spt6-1004* mutant, 284 (5.4%) were upregulated, and 1,133 (21.5%) were not significantly changed. Furthermore, the TFIIB ChIP-nexus signal also decreased for most genic promoters (Figure 2D), suggesting that the changes in the *spt6-1004* mutant are caused by changes in initiation, rather than a post-initiation step. We verified the decrease over the genic promoter of two genes by ChIP-qPCR of TFIIB (Figure 6A). Thus, results from assaying both transcription initiation and PIC levels showed that Spt6 plays a global role in the regulation of genic promoters.

**Figure 6.**
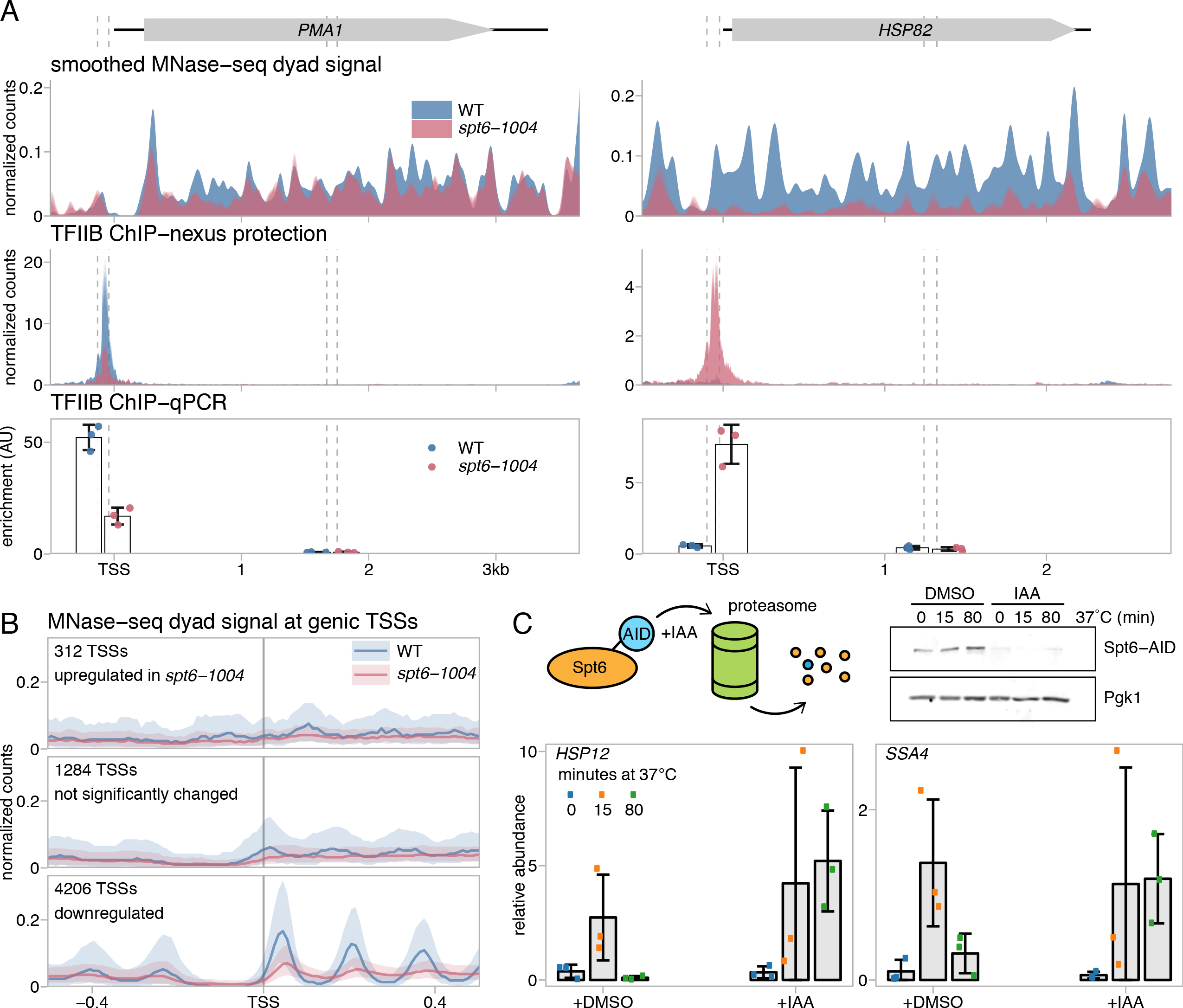
Spt6 function is necessary to control genic transcription. **(A)** ChIP-nexus and ChIP-qPCR measurements of TFIIB enrichment at the *PMA1* and *HSP82* genes in wild-type and spt6-1004 strains, plotted as in 2B. For the ChIP-qPCR analysis, the mean and standard deviation area plotted for three experiments. **(B)** The average MNase-seq dyad signal at genic TSSs in wild-type and *spt6-1004* strains, grouped by the differential expression status of the TSS. The solid line and shading represent the median and inter-quartile range. **(C)** RT-qPCR analysis of *HSP12* and *SSA4* RNA levels, testing the effects of temperature shift and Spt6 depletion. The top panel on the left shows a diagram of auxin-dependent degradation system used to deplete Spt6 and on the right shows a Western measuring the level of Spt6 protein, with and without depletion. The bottom panels show the RNA levels for *HSP12* and *SSA4* at times after a temperature shift from 30°C to 37°C. In these experiments, either DMSO (left side of each graph) or IAA (right side) were added 30 minutes before the zero time point. Plotted are the mean and standard deviation for three experiments, normalized to *SNR190* RNA.

To see whether promoter chromatin architecture might contribute to the differential regulation of genes by Spt6, we examined our MNase-seq data for the genic TSSs downregulated, upregulated, and not significantly changed in *spt6-1004.* Interestingly, each group has a distinct nucleosome profile (Figure 6B). Genes that are downregulated in *spt6-1004* and therefore require Spt6 for normal initiation have the expected wild-type profile of an NDR upstream of a strong +1 nucleosome peak. In the *spt6-1004* mutant, the MNase profile of these genes reflects the changes expected from the metagene MNase profile in Figure 4A, with a slightly shallower NDR and reduced +1 nucleosome occupancy (Figure 6B). In contrast, genes that are upregulated in *spt6-1004* and are therefore normally repressed by Spt6 have, on average, neither a detectable NDR nor a +1 nucleosome peak in either wild-type or *spt6-1004*. Finally, the genes not significantly affected in *spt6-1004* have a third nucleosome pattern which seems to be between the other two classes of genes. Thus, the three classes of genes differentially regulated by Spt6 have distinct chromatin architectures over their promoters.

Our analysis shows that the group of genes that are strongly repressed by Spt6 includes several that are normally induced by heat shock. To understand how Spt6 regulates this class of gene, we tested whether the induction of two genes, *SSA4* (Werner-Washburne et al., 1987) and *HSP12* (Praekelt and Meacock, 1990), require only the depletion of Spt6 or whether the induction also requires the temperature shift used to deplete Spt6 in the *spt6-1004* mutant. To separate the effects of Spt6 depletion and temperature shift, we used an auxin-inducible degron system (Nishimura et al., 2009) to deplete Spt6 in the absence of a temperature shift. We then measured RNA levels by RT-qPCR, independently varying Spt6 depletion and temperature shift. Our results (Figures 6D) show that both genes were induced only after a shift to 37°C, independently of whether Spt6 was depleted (see 15 minute time point). However, at 80 minutes after the temperature shift, at a time when adaptation to heat shock has normally occurred, RNA levels were still high when Spt6 was depleted. These results show that Spt6 is required for the repression of some heat shock-induced genes during adaptation after the temperature shift, consistent with a previously described function for the histone chaperone Spt16 (Jensen et al., 2008; Rowley et al., 1991) and with a role for Spt6 in repressing genes following their induction by carbon or phosphate starvation (Adkins and Tyler, 2006).

## DISCUSSION

In this work, we have integrated multiple quantitative genomic approaches to study the conserved transcriptional regulator Spt6 in *S. cerevisiae*, leading to new insights into Spt6 function and into the potential for expression of alternative transcripts. Our results have shown, for the first time on a genomic scale, that the thousands of intragenic and antisense transcripts produced in an *spt6* mutant are due to new transcription initiation from RNAPII transcriptional promoters. In addition, we identified sequence motifs at intragenic promoters that are also found at canonical promoters indicating that promoter-like sites exist broadly within genes and are normally maintained in a repressed state by Spt6. Furthermore, we showed that Spt6 plays a genome-wide role in the regulation of initiation from genic promoters. Together, these results demonstrate that Spt6 plays a critical role in determining the specificity of transcription initiation in vivo.

Our results support the idea that activation of intragenic promoters in an *spt6-1004* mutant is the consequence of nucleosome loss over regions that share some sequence characteristics of canonical promoters. These include the consensus sequence for initiation (Malabat et al., 2015; Zhang and Dietrich, 2005), a tendency to be more AT-rich, enrichment for TATA elements as previously described (Cheung et al., 2008; Uwimana et al., 2017), and enrichment for some transcription factor binding sites. In addition, intragenic initiation tends to occur in chromatin regions that are offset from nucleosome dyads in wild-type cells and that become nucleosome-depleted in an *spt6-1004* mutant.

The mechanism by which Spt6 normally represses thousands of intragenic promoters is uncertain. One study showed that Spt6 depletion allows ectopic localization of histone Htz1 in coding regions, suggesting that Spt6 represses intragenic promoters by excluding Htz1 (Jeronimo et al., 2015). However, our analysis suggests that the intragenic promoters that we have identified are not significantly enriched for the ectopic Htz1 locations previously found (data not shown). As Spt6 is also required for the recruitment of other proteins to transcribed chromatin, including the histone chaperone Spt2 (Chen et al., 2015; Nourani et al., 2006), as well as for histone H3 K36 methylation (Carrozza et al., 2005; Chu et al., 2006; Youdell et al., 2008), there are likely many aspects of Spt6 function that contribute to the repression of intragenic promoters. A recent study showed that antisense promoters have a distinctive pattern of histone modifications compared to sense promoters (Murray et al., 2015). It would be of interest to understand if this was also the case for intragenic promoters.

Our work has revealed that Spt6 is required for a normal level of transcription initiation from over 4,000 genic promoters. As Spt6 is primarily associated with transcribed regions (DeGennaro et al., 2013; Ivanovska et al., 2011; Mayer et al., 2010) and it has been shown to enhance the rate of elongation (Ardehali et al., 2009; Endoh et al., 2004), it was unexpected to discover that it regulates this class of initiation. We suggest that Spt6 regulates genic promoters indirectly, by controlling the total number of active promoters. In a wild-type yeast cell during growth in rich medium, there are ~5,000 expressed promoters and ~4,000-5,000 copies of most PIC proteins, including TFIIB (Ho et al., 2018). In contrast, in an *spt6-1004* mutant, there is a large increase in the number of active promoters, driving over 13,000 TSSs. Given that there is a decreased level of TFIIB in the *spt6-1004* mutant (~70% of wild-type levels), we suggest that the approximately three-fold increase in the number of TSSs results in a competition for a limited supply of PIC components, resulting in decreased expression from genic promoters. In support of this, our results show that in wild type there is a large difference in average expression levels between different classes of TSSs, while in the *spt6-1004* mutant, the differences in the expression levels between the classes are greatly diminished (Figure 1D), as if, in the mutant, all promoters have an approximately equal opportunity to recruit PICs.

Past studies of *spt6-1004* suggested that intragenic transcripts may encode functional information that is used in certain conditions (Cheung et al., 2008). Consistent with this idea, substantial evidence has emerged over the past few years that intragenic promoters occur throughout eukaryotes and that many are activated under particular growth conditions to carry out important functions. In addition to yeast, intragenic transcription occurs in mammalian cells in a widespread fashion under certain conditions (Carvalho et al., 2013; Muratani et al., 2014).

Furthermore, intragenic transcripts can encode N-terminally truncated proteins that have distinct functions compared to their full-length counterparts, including in oncogenes (Wiesner et al., 2015), during stress response (Tamarkin-Ben-Harush et al., 2017), and in the p53 family (Engelmann and Putzer, 2014; Wilhelm et al., 2010). In yeast and plants, functional N-terminally truncated proteins encoded by intragenic transcripts have also been demonstrated (Gammie et al., 1999; MacDiarmid et al., 2016; McKnight et al., 2014; Ushijima et al., 2017; Zhou et al., 2017). For two of the yeast genes that encode functional intragenic transcripts, *ASE1* and *KAR4*, we also observed intragenic initiation in *spt6-1004*. However, not all intragenic promoters that have been identified are active in *spt6-1004*. For example, a recent study showed that Gcn4 activates transcription from many intragenic sites (Rawal et al., 2018) and most of those are not activated in an *spt6-1004* mutant. In addition to encoding N-terminally truncated proteins, intragenic promoters can play other types of regulatory roles, such as interference with normal gene expression (Kim et al., 2017; Xie et al., 2011). The continued analysis of the regulation and function of intragenic transcription will likely lead to new insights into the flexibility of genomes in encoding functional information.

## SUPPLEMENTAL INFORMATION

Supplemental information includes four figures and two tables and can be found with this article online.

## AUTHOR CONTRIBUTIONS

S.M.D., O.V., M.M., L.S.C., and F.W. designed the experiments; S.M.D. performed the TSS-seq and ChIP-nexus experiments; O.V. performed the MNase experiments; M.M. performed the NET-seq experiments; D.S. performed the single gene ChIP, Western blots, and RT-qPCR experiments; J.C. performed and interpreted all of the bioinformatic analysis of the TSS-seq, ChIP-nexus, MNase-seq, and NET-seq datasets with input from S.M.D., L.S.C., and F.W.; S.M.D. and F.W. wrote the manuscript with feedback from all authors.

## ACKNOWLEDGMENTS

We thank Josh Arribere and Wendy Gilbert for critical advice on adapting TSS-seq from TL-seq; Burak Alver, Peter Park, and Julia di Iulio for bioinformatics support; Kevin Harlen, Ameet Shetty, and Rajaraman Gopalakrishnan for advice and discussions; Mary Couvillion and Blake Tye for helpful comments on the manuscript; and Natalia Reim for providing yeast strain FY3122. Part of this research was conducted on the O2 High Performance Computer Cluster supported by the Research Computing Group at Harvard Medical School. This work was supported by an American Cancer Society Fellowship to S.M.D.; NIH fellowship F32GM119291 to O.V., NIH grant R01HG007173 to L.S.C., and NIH grant R01GM032967 to F.W.

## CONTACT FOR REAGENT AND RESOURCE SHARING

Correspondence and requests for materials should be addressed to Fred Winston.

## EXPERIMENTAL MODEL AND SUBJECT DETAILS

Strains used in this study are listed in Table S1. All strains were constructed by standard procedures, using either yeast transformation or crosses. All oligonucleotides used for PCR are listed in Table S2. The *spt6-1004* temperature-sensitive mutant and wild-type strains were grown as previously described (Cheung et al., 2008): cells were grown in YPD at 30°C to a concentration of approximately 1 × 10^7^ cells/ml (OD600=0.6), at which point an equal volume of YPD medium pre-warmed to 44°C was added, and the cultures were shifted to 37°C for an additional 80 minutes.

## METHOD DETAILS

### Transcription start site sequencing

Yeast strains FY2180 and FY2181 were grown in 100 ml cultures at 30°C and shifted to 37°C as described above. After determining the cell concentration using a hemacytometer, *S. pombe* cells (strain FWP10) were added to each culture at a level of 10%, to be used for spike-in normalization. Total RNA was isolated as previously described (Ausubel, 1991). Poly(A)-enriched RNA was isolated from 300 mg of total RNA with 300 ml of Dynabeads oligo(dT)25 (Invitrogen), using the manufacturer’s instructions and eluted in water. Prior to each subsequent step of library construction, RNA samples were heat denatured at 80°C for two minutes and rapidly cooled on ice, followed by addition of 40 U of RNasin (Promega). Between each enzymatic reaction, samples were purified using an RNA binding column (Zymo Research). Ten to fifteen mg of poly(A) RNA was dephosphorylated with 30 units of calf intestinal phosphatase (CIP; NEB) for one hour at 37°C. CIP was removed from the reaction by heat inactivation followed by phenol extraction, and traces of phenol were removed using the above-mentioned RNA column. The m7GpppN cap was then cleaved from the RNA with 12.5 units of CapClip (CELLSCRIPT) for one hour at 37°C and the decapped RNA, containing a 5’ monophosphate, was ligated to 25 pmoles of a DNA/RNA chimeric linker (oSDAP4; Table S2) containing a randomized RNA linker sequence of six nucleotides at the 3’ end and a 5’ biotin moiety in a 10 ml reaction with 20 units of T4 RNA ligase 1 (NEB) and 2 mM ATP. Ligation products were column purified as before and eluted into fragmentation buffer (Ingolia et al., 2009) calibrated to enrich for 90-120 nucleotide oligomers. Fragmented RNA was then size selected and purified from a 10% acrylamide urea gel (Invitrogen). PNK removal of the 3’ phosphate group and 3’-end ligation of the RNA to a random linker pool (Mayer et al., 2015) was done as previously described (Couvillion and Churchman, 2017), except after ligation the biotinylated RNA was affinity purified with 10 ml of Dynabeads M-270 streptavidin (Invitrogen) using the manufactures instructions. Bead-bound RNA was eluted into 50 ml of elution buffer (0.1% SDS, 10 mM Tris 7.5) at 90°C for 5 minutes, and reverse transcribed with 3 pmoles of RT primer (oSMDRT2; Table S2) by heating for 5 min at 65°C, with 200 units SSIII Reverse Transcriptase (Invitrogen) at 48°C for 45 minutes. The cDNA was gel purified as above, and PCR amplified for 10-14 cycles using previously described indexing and sequencing primers for Illumina sequencing (Couvillion and Churchman, 2017).

### ChIP-qPCR and ChIP-nexus

For TFIIB studies, yeast strains FY3126 and FY3127 were grown in YPD at 30°C and then shifted to 37°C as described above. The cultures were cooled to 25°C using pre-chilled medium at 4°C before cross-linking in 1% formaldehyde while shaking at 25°C for 30 minutes, followed by quenching in 125mM glycine at 25°C for 10 minutes. For Spt6 and Rpb1 ChIP-nexus, strain FY3128 was grown without the temperature shift. Chromatin was extracted using standard methods (DeGennaro et al., 2013) and sheared in a QSONICA sonicating water bath. For ChIP-qPCR spike-in normalization, each *S. cerevisiae* chromatin sample was mixed with 50% *S. pombe* chromatin (strain FWP561) by mass for TFIIB ChIP and 30% by mass for histone H3 ChIP. Chromatin precipitations were performed overnight at 4°C with 4 μg of anti-H3 (ab1791; Abcam) per 300 μg of chromatin or 20 μl of Pan Mouse IgG Dynabeads (Invitrogen) per 500 μg of chromatin. Real-time qPCR was performed as previously described (DeGennaro et al., 2013) using primer pairs listed in Table S2.

Each ChIP-nexus library used 2.5-3 mg of *S. cerevisiae* chromatin containing 5% *S. pombe* chromatin added by mass for downstream spike-in normalization between samples (see ChIP-nexus library processing section below). To generate sequencing libraries for TFIIB and Spt6 bearing TAP tags, chromatin was affinity purified using 100 ml Pan Mouse IgG Dynabeads (Invitrogen). For RNAPII (Rbp1) libraries, chromatin was immunoprecipitated with 40 mg of 8WG16 antibody (BioLegend) that was pre-bound to 100 ml of ProteinG Dynabeads (Invitrogen). Library constructions for Illumina sequencing were performed essentially as previously described (He et al., 2015), except buffers were optimized for yeast: Buffer A (10 mM TE, 0.1% Triton X), Buffer B (50 mM HEPES.KOH pH 7.4, 140 mM NaCl, 1 mM EDTA, 1% Triton-X, 0.1% sodium deoxycholate), Buffer C (Buffer B with 250 mM NaCl), Buffer D (10 mM Tris pH 7.5, 250 mM LiCl, 10 mM EDTA, 0.5% IGEPAL CA-360, 0.1% sodium deoxycholate).

### MNase-seq

MNase digestion was performed as previously described (Rando, 2010) with some modifications, using strains FY87 and FY3125. Cultures of 500 ml were grown in YPD at 30°C, then shifted to 37°C as described above. At a density of approximately 1 × 10^7^ cells/ml (OD600 = 0.5), crosslinked using 2% formaldehyde for 30 minutes and then treated for 10 minutes with 125 mM glycine before collecting an equal number of cells for each strain. The cells were resuspended in 40 ml of sorbitol buffer (1 M sorbitol, 50 mM Tris pH7.4, 10 mM β-mercaptoethanol) and incubated for 30 minutes at 30°C with 10 mg of zymolase 100T (US Biological) per gram of cells. Spheroplasting efficiency was assessed by microscopy and was more than 95% of total cells. The spheroplasts were collected and resuspended in NP buffer (1 M sorbitol, 50 mM sodium chloride, 10 mM Tris pH 7.4, 5 mM magnesium chloride, 1 mM calcium chloride, 0.075% NP-40, 1 mM β-mercaptoethanol, 500 μM spermidine). Micrococcal nuclease (MNase; Sigma) was dissolved in Ex50 buffer (10 mM Hepes pH 7.6, 50 mM sodium chloride, 1.5 mM magnesium chloride, 0.5 mM EGTA, 10% glycerol, 1mM dithiothreitol, 0.2 mM phenylmethylsulfonyl fluoride)prepared to produce 500 units per 840 μl stock as recommended by the manufacturer. The spheroplasts were divided into aliquots and incubated for 20 minutes at 37°C with increasing amounts of MNase, ranging from 2 to 15 μl of the stock. Digestion was stopped by addition of stop buffer (5% SDS, 100 mM EDTA), samples were subjected to proteinase K digestion and reverse-crosslinking at 65°C overnight, followed by DNA purification. The efficiency of MNase digestion was quantified using DNA fragment size analysis (Agilent Bioanalyzer) to establish an MNase titration curve for each strain. The MNase concentrations which yield approximately 80% mononucleosomal DNA were selected for the library construction. The samples were mixed with the MNase-digested spike-in DNA from *S. pombe* based on the original cell count (100 ng of spike-in DNA per MNase digested DNA from 7 × 10^8^ *S. cerevisiae* cells). Mononucleosomal DNA was purified using size-selected gel extraction. The sequencing libraries were constructed as described before (DeGennaro et al., 2013).

### NET-seq

NET-seq was performed on strains grown at both 30°C and 37°C. Strains FY2912 and FY2913 were grown at 30°C, the cultures were split and half was shifted to 37°C as described above. NET-seq was performed as previously described (Churchman and Weissman, 2011).

### Western blotting

To measure FLAG-Spt6 and TFIIB-TAP protein levels, strains FY3126 and FY3127 were grown with the 37°C temperature shift as described above. Prior to pelleting the cells, strain FY2354 expressing *DST1-MYC* was added to each culture at 50% concentration by cell number used for spike-in normalization. Cell extracts were made by bead beating in LB-140 buffer (50 mM HEPES.KOH pH 7.4 140 mM NaCl 1 mM EDTA 1% TritonX-100 0.1% NaDeoxycholate0.1% SDS) along with protease inhibitors (1mM phenylmethylsulfonyl fluoride, 2 μg/mL leupeptin, 2 μg/mL pepstatin, 0.4 mM dithiothreitol), and SDS-PAGE gels were loaded by mass. For protein detection, primary antibodies used were anti-FLAG diluted 1:5000 (clone M2; SIGMA), anti-Protein A diluted 1:1500 (clone SPA-27; SIGMA), anti-cMyc diluted 1:1000 (clone A-14 Santa Cruz), anti-PGK1 diluted 1:20000 (clone 22C5D8; Invitrogen) and anti-V5 diluted 1:2000 (clone R960-25; Invitrogen). Secondary detection used anti-mouse and anti-rabbit IR-dye-coupled antibodies from Li-Cor Biosciences. Protein bands were detected using the Li-Cor Aerius and intensities were quantified by measuring their integrated density with Adobe Photoshop Extended version 19.1.4.

### Auxin induced degradation

Yeast strain FY3122 was grown in YPD at 30°C to a concentration of approximately 1 × 10^7^ cells/ml (OD600=0.6), at which point cells were treated with 25 μM 3-Indoleacetic acid (IAA; SIGMA) for 30 minutes prior to shifting to 37°C as described above. Samples were taken for Western (see above) and RT-qPCR analysis at the indicated timepoints described in the text. RT-qPCR was done as previously described (DeGennaro et al., 2013). Primer pairs for *SSA4* and *HSP82* genes were as previously published (Anandhakumar et al., 2016) and listed in Table S2.

### Data management

All data analyses were managed using the Snakemake workflow management system (Koster and Rahmann, 2012), and are available at github.com/winston-lab.

### TSS-seq library processing

Removal of adapter sequences from the 3’ end of the read and 3’ quality trimming were performed using cutadapt (Martin 2017). The random hexamer molecular barcode on the 5’ end of the read was then removed and processed using a custom Python script (Mayer et al., 2015). Reads were aligned to the combined *S. cerevisiae* and *S. pombe* reference genomes using Tophat2 without a reference transcriptome (Kim et al., 2013), and uniquely mapping reads were selected using SAMtools (Li et al., 2009). Reads mapping to the same location as another read with the same molecular barcode were identified as PCR duplicates and removed using a custom Python script (Mayer et al., 2015). Coverage of the 5’-most base, corresponding to the TSS, was extracted using bedtools genomecov (Quinlan and Hall, 2010) and normalized to the total number of reads uniquely mapping to the *S. pombe* genome. Quality statistics of raw, cleaned, non-aligning, and uniquely aligning non-duplicate reads were assessed using FastQC (Andrews, 2014).

### TSS-seq peak calling

TSS-seq data for a single TSS tends to occur as a group of highly-correlated signals over a window of nucleotides, rather than at a single nucleotide. Therefore, for identification of TSSs and quantification for analyses such as differential expression, it is necessary to perform peak-calling. TSS-seq peak calling was performed using a 1-D watershed segmentation algorithm, followed by filtering for reproducibility by the Irreproducible Discovery Rate (IDR) method (Boleu et al., 2015; Li et al., 2011). First, a smoothed version of the TSS-seq coverage was generated for each sample using adaptive two-stage kernel density estimation with a discretized Gaussian kernel (pilot bandwidth = 10 nt, bandwidth = 10 nt, *α* = 0.2). The adaptive kernel adjusts the kernel bandwidth to be smaller in regions of high signal density and larger in regions of lower signal density (Silverman, 1986), allowing the smoother to better accommodate both ‘sharp’ TSSs where the signal is distributed over a relatively small window as well as ‘broad’ TSSs where the signal is more dispersed. Following smoothing, an initial set of peaks is formed by assigning all nonzero signal in the original, unsmoothed coverage to the nearest local maximum of the smoothed coverage, and taking the minimum and maximum genomic coordinate of the original coverage as the peak boundaries for each local maximum of the smoothed coverage. Peaks are then trimmed to the smallest genomic window that includes 95% of the original coverage, and the probability of the peak being generated by noise is estimated by a Poisson model where *λ*, the expected coverage, is the maximum of the expected coverage over the chromosome and the expected coverage in the 2000 nt window upstream of the peak (as for the ChIP-seq peak caller MACS (Zhang et al., 2008b)). Finally, peaks are ranked by their significance under the Poisson model, and a final list of peaks for each condition is generated using the IDR method (`IDR = 0.1`) (Boleu et al., 2015; Li et al., 2011).

### TSS-seq differential expression analysis

For TSS-seq differential expression, TSS-seq peak-calling was performed as described above for both *S. cerevisiae* and the *S. pombe* spike-ins. The read counts for each peak in each condition were used as the input to differential expression analysis by DESeq2 (Love et al., 2014), with the alternative hypothesis |log_2_(fold - change)| > 1.5 and a false discovery rate of 0.1. To normalize by spike-in, the size factors of the *S. pombe* spike-in counts were used as the size factors for *S. cerevisiae*, although we note that due to the median of ratios normalization method used in DESeq2, the major TSS-seq results of this work are still observed when the *S. cerevisiae* size factors are used.

### ChIP-nexus library processing

Filtering for reads containing the constant region of the adapter on the 5’ end of the read, 3’ adapter removal and 3’ quality trimming were performed using cutadapt (Martin, 2017). The random pentamer molecular barcode on the 5’ end of the read was then removed and processed using a modified custom Python script (Mayer et al., 2015). Reads were aligned to the combined *S. cerevisiae* and *S. pombe* genomes using Bowtie2 (Langmead and Salzberg, 2012), and uniquely mapping reads were selected using SAMtools (Li et al., 2009). Reads mapping to the same location as another read with the same molecular barcode were identified as PCR duplicates and removed using a custom Python script (Mayer et al., 2015). Coverage of the 5’-most base, corresponding to the point of crosslinking, was extracted using bedtools genomecov (Quinlan and Hall, 2010). The median fragment size estimated by MACS2 (Zhang et al., 2008b) over all samples was used to generate coverage of factor protection and fragment midpoints, by extending reads to the fragment size, or by shifting reads by half the fragment size, respectively. Coverage was normalized to the total number of reads uniquely mapping to *S. cerevisiae.* Quality statistics of raw, cleaned, non-aligning, and uniquely aligning non-duplicate reads were assessed using FastQC (Andrews, 2014).

### TFIIB ChIP-nexus peak-calling

TFIIB ChIP-nexus peak calling was performed using MACS2 (Zhang et al., 2008a), using 160 bp for the model-building bandwidth, 1000bp as the size of the large local region used to model expected counts, and the default false discovery rate of 0.05. Reads mapping to the same base were kept since PCR duplicates were filtered out using the molecular barcode. MACS2 was chosen over several ChIP-nexus and ChIP-exo specific peak calling tools because the specialized tools tended to split each TFIIB peak into multiple subpeaks, likely due to the multiple crosslinking points of TFIIB to the DNA (Rhee and Pugh, 2012).

### Reannotation of *S. cerevisiae* TSSs using TSS-seq data

TSS-seq coverage from two replicates of a wild-type *S. cerevisiae* strain grown at 30°C in YPD (data not shown) was averaged and used to adjust the 5’ ends of an annotation file of major transcript isoforms based on TIF-seq data (Pelechano et al., 2013). The 5’ end of the original annotation was changed to the position of maximum TSS-seq signal in a window 250nt in each direction if the TSS-seq signal at that position was greater than the 95^th^ percentile of all non-zero TSS-seq signal.

### Classification of TSS-seq and TFIIB ChIP-nexus peaks into genomic categories

TSS-seq and TFIIB ChIP-nexus peaks were assigned to genomic categories based on their position relative to the transcript annotation described above and an annotation of all verified open reading frames (ORF) and blocked reading frames in *S. cerevisiae* (Crooks et al., 2004; Engel et al., 2014). First, genic regions were defined as follows: If a gene was present in both the transcript and ORF annotations, the genic region was defined as the interval (annotated TSS - 30 nucleotide, start codon]. If a gene was present in the transcript annotation but not the ORF annotation, the genic region was defined as the interval (annotated TSS-30nt, annotated TSS+30nt]. If a gene was present only in the ORF annotation, the genic region was defined as the interval (start codon-30nt, start codon]. For the purposes of peak classification, regions were considered overlapping if they had at least one base of overlap. Peaks were classified as genic if they overlapped a genic region on the same (TSS-seq) or either (TFIIB ChIP-nexus) strand. Peaks were classified as intragenic if they were not classified as a genic peak, and additionally overlapped an open or closed reading frame on the same (TSS) or either (TFIIB ChIP-nexus) strand. TSS-seq peaks were classified as antisense if they overlapped a transcript on the opposite strand. TSS-seq and TFIIB ChIP-nexus peaks were classified as intergenic if they did not overlap a transcript, reading frame, or genic region on either strand.

### TSS information content

TSS-seq alignments were pooled for all replicates in a condition, and the DNA sequence flanking the position of every read overlapping TSS-seq peaks of a particular genomic category was extracted using SAMtools (Li et al., 2009) and bedtools (Quinlan and Hall, 2010). The information content of the sequences was quantified with WebLogo (Crooks et al., 2004), with the zeroth-order Markov model of the *S. cerevisiae* genomic sequence as the background composition. Sequence logos were plotted with helper functions from ggseqlogo (Wagih, 2017).

### TFIIB ChIP-nexus differential binding analysis

For TFIIB ChIP-nexus differential binding analysis, TFIIB peaks were called as described above. A non-redundant list of peaks called in any condition was generated using bedtools, and the counts of fragment midpoints for each peak in each condition were used as the input to differential binding analysis by DESeq2 (Love et al., 2014), with the alternative hypothesis |log_2_(fold - change)| > 2 and a false discovery rate of 0.1. For estimation of changes in TFIIB binding upstream of TSS-seq peaks, TFIIB fragment midpoint counts were used as the input to differential binding analysis by DESeq2, using *S. cerevisiae* counts for size factors.

### NET-seq library processing

Removal of adapter sequences from the 3’ end of the read and 3’ quality trimming were performed using cutadapt (Martin, 2017). Reads were aligned to the *S. cerevisiae* genome using Tophat2 without a reference transcriptome (Kim et al., 2013), and uniquely mapping reads were selected using SAMtools (Li et al., 2009). Coverage of the 5’-most base of the read, corresponding to the 3’-most base of the nascent RNA and the active site of elongating RNA polymerase, was extracted using bedtools genomecov (Quinlan and Hall, 2010) and normalized to the total number of uniquely mapped reads. Quality statistics of raw, cleaned, non-aligning, and uniquely aligning reads were assessed using FastQC (Andrews, 2014).

### MNase-seq library processing

Paired-end reads were demultiplexed using fastq-multx (Aronesty, 2103), allowing one mismatch to the barcode. Read 2 barcode removal and 3’ quality trimming were performed with cutadapt (Martin, 2017). Reads were aligned to the combined *S. cerevisiae* and *S. pombe* genome using Bowtie 1 (Langmead et al., 2009), and correctly paired reads selected using SAMtools (Li et al., 2009). Coverage of nucleosome protection and nucleosome dyads were extracted using bedtools (Quinlan and Hall, 2010) and custom shell scripts to get the entire fragment or the midpoint of the fragment, respectively. Smoothed nucleosome dyad coverage was generated by smoothing dyad coverage with a Gaussian kernel of 20 bp bandwidth. Coverage was normalized to the total number of correctly paired *S. cerevisiae* fragments. Quality statistics of raw, cleaned, non-aligning, and correctly pairing reads were assessed using FastQC (Andrews, 2014).

### MNase-seq quantification

Quantifications of nucleosome occupancy, fuzziness, and position shifts were calculated using DANPOS2 (Chen et al., 2013) with the total counts in mutant libraries scaled by the mean observed spike-in percentage in the mutant libraries over the mean observed spike-in percentage in the wild-type libraries for spike-in normalization.

### Clustering of MNase-seq signal at *spt6-1004* intragenic TSSs

Spike-in normalized MNase-seq dyad signal in the window 150bp to either side of the summit of the 6059 intragenic TSS-seq peaks upregulated in *spt6-1004* over wild-type was binned by taking the mean signal in non-overlapping 5bp bins, and then averaged by taking the mean of two replicates *(spt6-1004)* or one experiment (wild-type). The wild-type and *spt6-1004* data were used as equally weighted 6059×60 input layers to a super-organizing map (SOM)(Wehrens and Buydens, 2007) trained using the input data to assign similar MNase-seq observations in 60-dimensional input space to similar nodes in a 2-dimensional (6×8) rectangular grid. The 48 ‘code vectors’ representing the typical MNase-seq pattern for each node were then clustered by agglomerative hierarchical clustering using sum of squares distance and Ward linkage. The resulting dendrogram was cut to produce the two clusters of MNase-seq signal shown in Figure 5. The choice to cut the dendrogram to produce two clusters was made because clusters created from deeper cuts tended to have nucleosome phasing patterns similar to the original two clusters. We note that the two clusters are stable under repeated training of the SOM with different random seeds. By chance, some random seeds will result in a third cluster which joins after the two major clusters have joined in the hierarchical clustering. However, this cluster is usually much smaller than the major clusters (<20 iTSSs) and can be grouped visually into one of the two major phasing patterns.

### Intragenic TSS position bias

As TSS-seq peaks are required to not overlap genic regions in order to be classified as intragenic, the expected distribution if intragenic TSSs were randomly distributed along the length of an ORF is not uniform. Therefore, the expected random distribution of intragenic TSSs was determined by taking all position of the ORF that the TSS could have taken and still been called intragenic. The random distribution was then compared to the observed distribution of intragenic starts by binning start locations to the nearest tenth of a percentage of relative distance along the ORF, and applying a permutation test on the chi-squared test statistic.

### Motif enrichment

FIMO (Grant et al., 2011) was used to search the *S. cerevisiae* genome for 3010 motifs from six databases (de Boer and Hughes, 2012; MacIsaac et al., 2006; Newburger and Bulyk, 2009; Pachkov et al., 2013; Teixeira et al., 2018; Zhu and Zhang, 1999). The zeroth-order Markov model of the *S. cerevisiae* genome sequence was used as a background model, with a p-value cutoff of 1e-5. For determining the enrichment of motif sites upstream of TSSs, the regions extending 200 base pairs upstream of TSS summits were taken and merged if they were overlapping. Motifs were considered to be present in a region if the entire motif was overlapping the region. The frequency of motif occurrences in the regions of interest was compared to the frequency of occurrences in the regions upstream of 6000 randomly chosen locations, using Fisher’s exact test.

### Enrichment of TATA boxes

Enrichment of TATA boxes was tested as for the other motifs described above, except for the following differences: First, the query motif used was TATAWAWR, where the ambiguous bases are equiprobable. Second, the p-value was 6e-4, chosen because it was the threshold required for only exact matches to be returned. Third, the TATA motif was required to be on the sense strand relative to the TSS in order to be counted as a match.

## QUANTIFICATION AND STATISTICAL ANALYSIS

Quantification and statistical tests employed for each experiment are described in the figure legends or in the methods section.

## DATA AND SOFTWARE AVAILABILITY

The raw sequencing data reported in this paper has been deposited at the NCBI Gene Expression Ominbus, accession number GSE115775.

